# Pharmacological Activation of NRF2 by Omaveloxolone Upregulates NRF2-Target Proteins in SMA Type I Human Fibroblasts

**DOI:** 10.64898/2026.03.17.712434

**Authors:** Sofia Vrettou, Sebastian Zetzsche, Brunhilde Wirth

**Author notes:** Corresponding author: Sofia Vrettou.

## Abstract

Spinal muscular atrophy (SMA) is caused by loss of SMN protein and is increasingly recognized as a multisystem disorder involving molecular pathology beyond motor neurons. Recently, we identified NRF2-KEAP1 signaling as dysregulated in SMA mice. Because NRF2 coordinates transcriptional programs that maintain cellular redox homeostasis and adaptive stress responses, we investigated whether NRF2 signaling is similarly altered in SMA type I patient-derived fibroblasts and whether it can be pharmacologically engaged. Compared with control fibroblasts, SMA fibroblasts displayed reduced basal expression of NRF2 target proteins, including NQO1 and xCT (SLC7A11), along with decreased levels of PGC1_α_. Omaveloxolone (OMAV), a pharmacological NRF2 activator approved for the treatment of Friedreich’s ataxia, increased cell viability and upregulated NRF2 target proteins in both control and SMA fibroblasts. Notably, OMAV produced a modest increase in SMN protein abundance and PGC1_α_ levels selectively in SMA cells. Together, these findings support diminished NRF2 pathway output as a feature of SMA fibroblasts and demonstrate that OMAV induces NRF2 target proteins in this human SMA cellular model, consistent with enhanced cytoprotective signaling.

**Graphical abstract:** 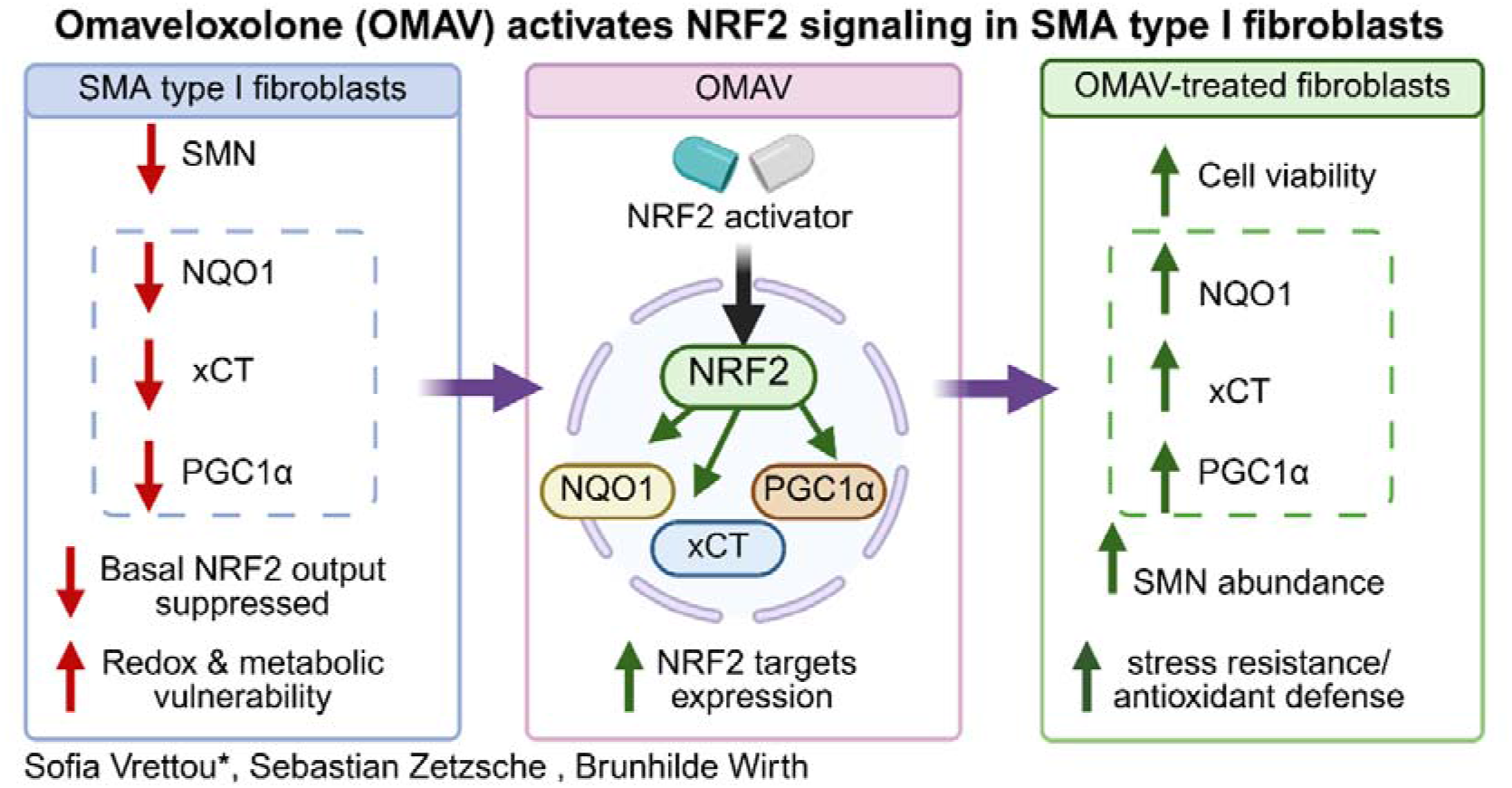

## Introduction

Spinal muscular atrophy (SMA) is an autosomal recessive neuromuscular disorder caused by loss-of-function mutations in *SMN1*, resulting in insufficient levels of survival motor neuron (SMN) protein [1, 2]. Disease severity is modified by *SMN2* copy number, which partially compensates for *SMN1* loss through limited production of full-length SMN [3]. SMN is ubiquitously present and participates in multiple fundamental cellular processes, including small nuclear ribonucleoprotein (snRNP) assembly and RNA metabolism [4]. Although motor neuron degeneration defines the clinical phenotype, accumulating evidence supports broader multisystem involvement, including metabolic dysfunction and redox imbalance in peripheral tissues [5, 6]. Consistent with this concept, our most recent multi-organ analyses in SMA mouse models have revealed organ-specific redox imbalance that is partially normalized following SMN restoration [7]. Moreover, we have shown that the NRF2-KEAP1 pathway is dysregulated in this system [8].

The NRF2-KEAP1 pathway is a central cytoprotective axis regulating detoxification enzymes, glutathione metabolism, and oxidative stress adaptation [9, 10]. Canonical NRF2 targets include NAD(P)H quinone dehydrogenase 1 (NQO1) and the cysteine-glutamate antiporter SLC7A11 (xCT), which support redox buffering through quinone detoxification and cystine import for glutathione synthesis, respectively [11, 12]. In addition, mitochondrial dysfunction and oxidative stress have been reported in SMA models, suggesting that redox imbalance contributes to cellular vulnerability beyond the neuromuscular compartment [13-15].

## Results & Discussion

Patient-derived fibroblasts represent a useful cellular model to investigate systemic molecular alterations associated with SMN deficiency, despite not being a primary disease tissue. However, endogenous antioxidant signaling in human SMA patient-derived fibroblasts has not been fully defined.

Here, we first assessed whether pharmacological modulation of NRF2 signaling influences cellular viability. Control (*SMN1*: 2 copies; *SMN2*: 1-2 copies) and SMA type I (*SMN1*: 0 copies; *SMN2*: 2 copies) fibroblasts were treated with NRF2-modulating compounds including sulforaphane (SFN), dimethyl fumarate (DMF), and the antioxidant precursor N-acetylcysteine (NAC). Cells were exposed to two concentrations of each compound and cell viability was monitored using the MTT assay following daily compound administration (Figure 1A).

**FIGURE 1:**
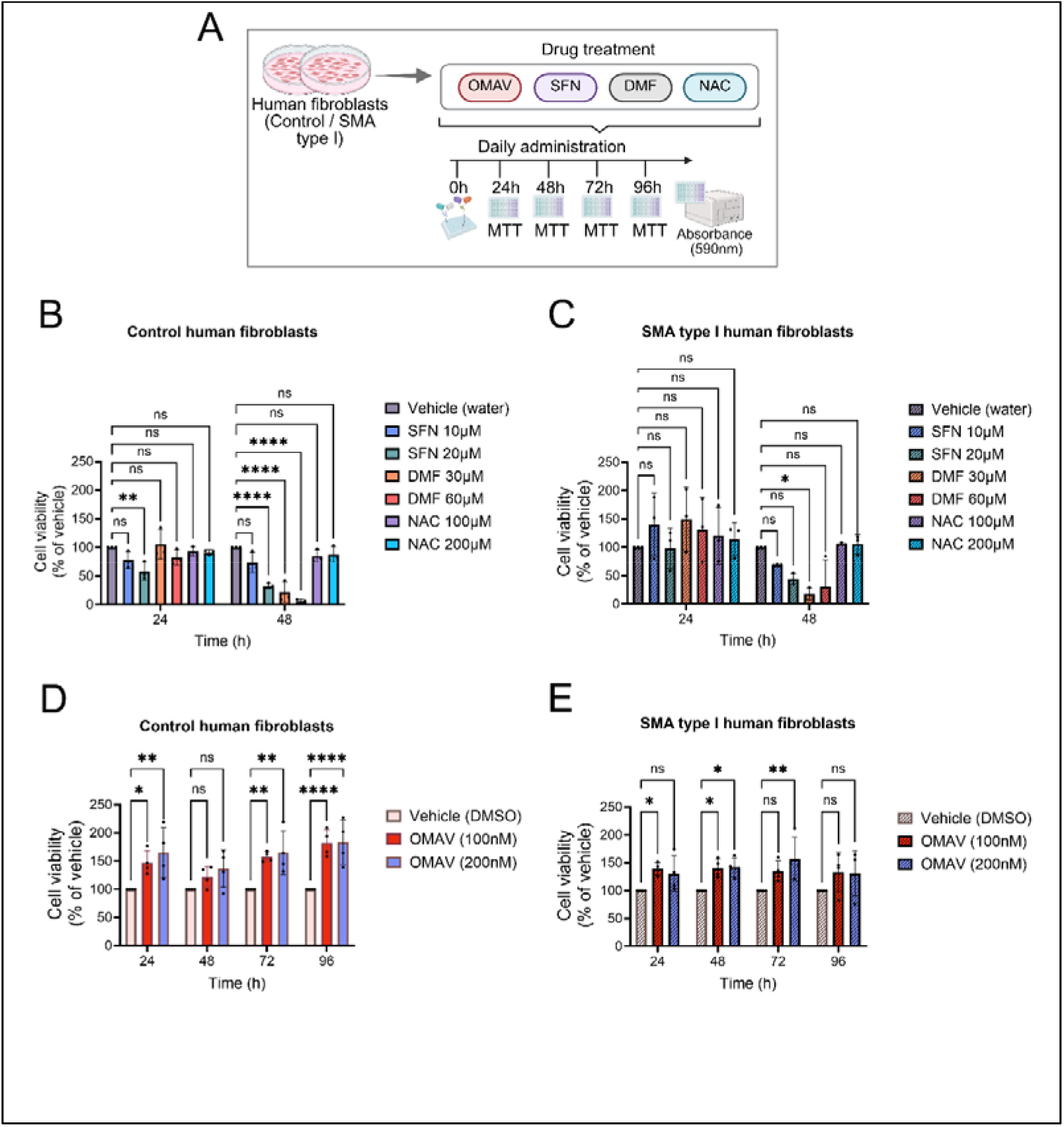
Pharmacological modulation of cell viability in control and SMA type I human fibroblasts. (A) Experimental design. Control and SMA type I human fibroblasts were treated daily with NRF2-modulating compounds: sulforaphane (SFN), dimethyl fumarate (DMF), omaveloxolone (OMAV) or the antioxidant precursor N-acetylcysteine (NAC), or the corresponding vehicles. Cell viability was assessed by MTT assay at the indicated time points (24 h, 48 h, 72 h and 96 h); absorbance was measured at 590 nm. (B-C) Compound screen in control (B) and SMA type I (C) fibroblasts treated with SFN (10 or 20 µM), DMF (30 or 60 µM), NAC (100 or 200 µM). Viability was measured at 24 and 48 h. SFN, DMF, and NAC were prepared in water, and cell viability values were normalized to the corresponding vehicle controls and expressed as % cell viability. (D-E) OMAV time course in control (D) and SMA type I (E) fibroblasts treated with OMAV (100 nM or 200 nM) or vehicle (DMSO) with daily refreshment. Viability was measured at 24 h, 48 h, 72 h, and 96 h, normalized to DMSO vehicle control, and expressed as % cell viability. The final DMSO concentration in culture was ∼0.001% (v/v). Data represent 3-4 independent fibroblast cell lines per genotype and are shown as mean ± SD. Statistical analysis was performed using one-way ANOVA followed by Dunnett’s multiple comparisons test within each timepoint. Statistical significance is indicated as: *p < 0.05; **p < 0.01; ****p < 0.0001; ns, not significant.

In control fibroblasts, higher concentrations of SFN and DMF reduced cell viability, whereas NAC did not significantly alter viability compared with vehicle-treated cells (Figure 1B). A similar pattern was observed in SMA type I fibroblasts. SFN and DMF decreased cell viability at higher doses, while NAC again showed no significant effect on cell survival (Figure 1C). These findings indicate that neither NRF2 activation by SFN or DMF nor antioxidant supplementation with NAC improved cellular viability under these experimental conditions.

We next examined the impact of omaveloxolone (OMAV), a potent pharmacological NRF2 activator approved for the treatment of Friedreich’s ataxia [16], on SMA cells. Treatment with OMAV (100 nM or 200 nM) increased cell viability in both control and SMA type I fibroblasts and was well tolerated at both concentrations (Figure 1D, E).

Because the increase in cell viability was already detectable at the 48-hour time point following daily OMAV administration, we selected this time window to examine OMAV-induced changes in protein expression. Cells were therefore treated with OMAV every 24 hours for two consecutive days prior to Western blot analysis (Figure 2).

**FIGURE 2:**
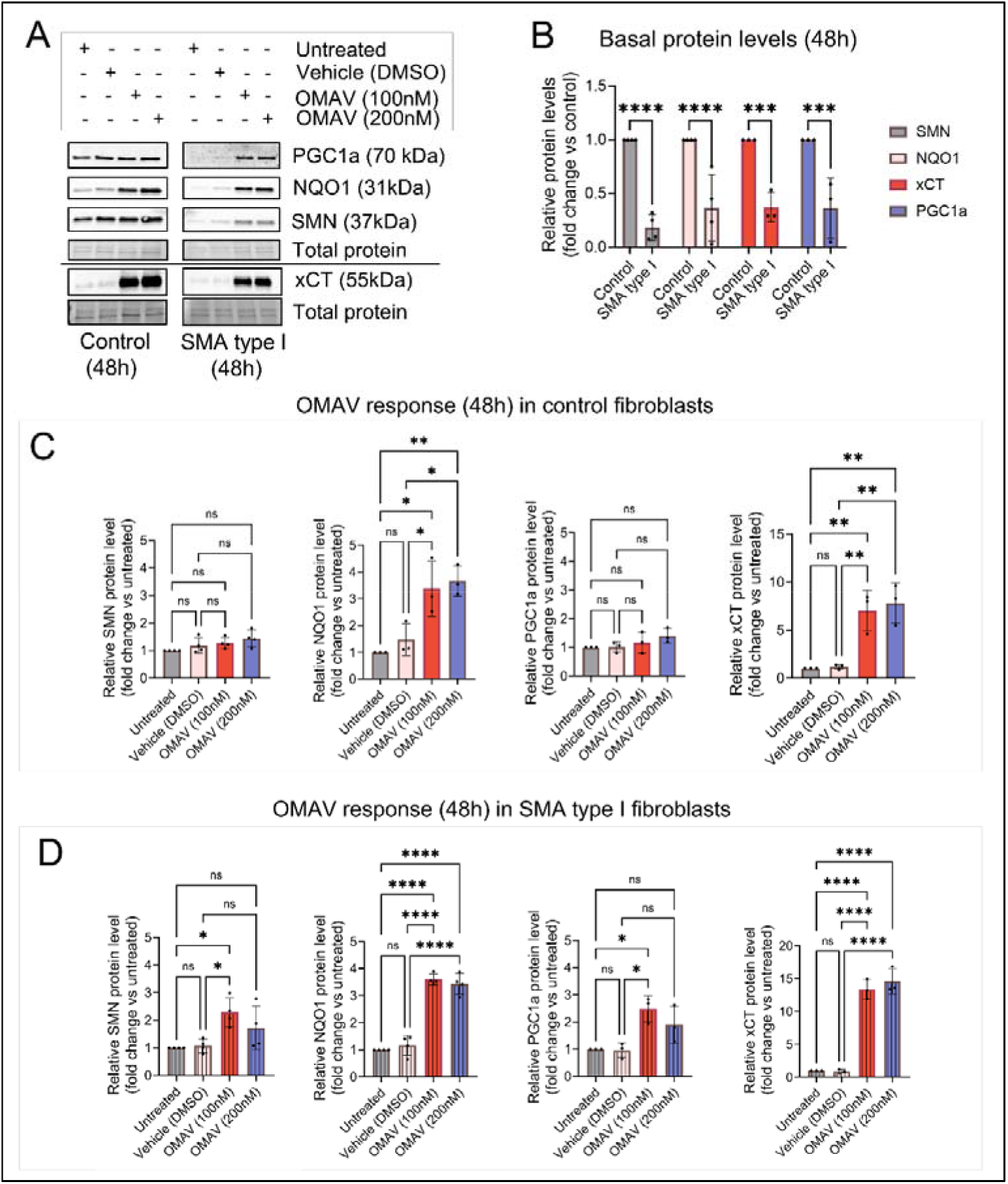
Reduced basal NRF2 target proteins in SMA type I fibroblasts and induction by omaveloxolone. (A) Representative Western blots from control and SMA type I fibroblasts treated for 48 h with omaveloxolone (OMAV; 100 nM or 200 nM), vehicle (DMSO), or left untreated. Membranes were probed with antibodies against SMN, NQO1, PGC1_α_ and xCT. Total protein staining was used as the loading control for normalization. (B) Basal protein levels (untreated) in control versus SMA type I fibroblasts, normalized to total protein and expressed relative to control (set to 1) for each target. Statistical analysis was performed using two-way ANOVA with Šídák’s multiple-comparisons test comparing control vs SMA for each protein target. (C-D) OMAV response in control (C) and SMA type I (D) fibroblasts treated with OMAV (100 nM or 200 nM). Protein abundance was normalized to total protein and expressed as fold change relative to the untreated condition (set to 1), because vehicle (DMSO) produced a mild increase in SMN in some cell lines. Statistical comparisons are shown for untreated vs all conditions and for vehicle (DMSO) vs all conditions, as indicated on the graphs. Panels C-D were analyzed by one-way ANOVA followed by Šídák’s multiple-comparisons test. Dots represent independent fibroblast cell lines and are presented as mean ± SD. Three control and three SMA type I cell lines were analyzed; one line pair was repeated in an independent experiment, yielding n = 4 for SMN and NQO1 and n = 3 for xCT and PGC1_α_. Statistical significance is indicated as: *p < 0.05; **p < 0.01; ***p < 0.001; ****p < 0.0001; ns, not significant.

Western blot analysis revealed reduced basal levels of NRF2 target proteins in SMA fibroblasts compared with controls (Figure 2A, B), including NQO1 and xCT. SMA fibroblasts also exhibited decreased levels of peroxisome proliferator-activated receptor gamma coactivator-1 alpha (PGC1_α_) (Figure 2A, B), a key regulator of mitochondrial metabolism and biogenesis [17]. Although PGC1_α_ is not a canonical NRF2 target, functional crosstalk between NRF2 signaling and PGC1_α_-dependent metabolic programs has been reported [18], linking redox homeostasis with mitochondrial metabolic regulation. The reduction of PGC1_α_ observed here is therefore consistent with growing evidence that SMN deficiency is associated with metabolic and mitochondrial perturbations in patient-derived fibroblasts, including recent reports of mitochondrial dysfunction even in fibroblasts from SMA carriers [19].

OMAV increased NQO1 and xCT levels in both control and SMA fibroblasts, consistent with induction of NRF2 target proteins (Figure 2C, D). In control fibroblasts, PGC1_α_ and SMN remained unchanged (Figure 2A, C). In contrast, in SMA fibroblasts, OMAV induced a more prominent increase in NQO1 and xCT relative to untreated cells and significantly increased both PGC1_α_ and SMN at the 100 nM dose (Figure 2A, D). These findings indicate that OMAV activates NRF2-associated target expression in both control and SMA fibroblasts, while selectively increasing PGC1_α_ and SMN in SMA cells.

The mechanism underlying the modest increase of SMN in SMA fibroblasts remains unclear. Previous studies have suggested that cellular stress and redox imbalance can influence SMN stability and turnover [14, 20], linking SMN homeostasis to proteostasis and metabolic state. Given NRF2’s broad cytoprotective roles in redox control and metabolic adaptation [10], NRF2 activation could indirectly affect SMN protein stability or translation. Alternatively, improved cellular homeostasis following NRF2 activation could create conditions that favor SMN protein accumulation in SMN-deficient cells. Elucidating the molecular relationship between NRF2 signaling and SMN regulation will require targeted mechanistic studies.

Collectively, our findings demonstrate that:

1. SMA type I human fibroblasts exhibit reduced basal expression of NRF2 target proteins, including NQO1 and xCT, together with decreased PGC1_α_.
2. Pharmacological activation of NRF2 signaling by OMAV increases expression of NRF2 target proteins in both control and SMA fibroblasts.
3. OMAV modestly increases SMN protein abundance in SMA fibroblasts.

Together, these results identify reduced NRF2 pathway output as a feature of SMN-deficient fibroblasts and show that pharmacological NRF2 activation enhances expression of cytoprotective target proteins in this cellular context. Further studies will be required to define the mechanistic relationship between NRF2 signaling and SMN regulation and to evaluate the potential translational implications of NRF2 activation in SMA.

## Ethics Statement

Informed written consent was obtained from individuals with SMA and controls according to the Declaration of Helsinki, and the study was approved by the ethics committee of the University Hospital of Cologne under the approval numbers 04-138.

## Author Contributions

**Sofia Vrettou:** Designed the work, performed all the experiments, analyzed the results and wrote the manuscript. **Sebastian Zetzsche:** Involved in cell culture experiments. **Brunhilde Wirth:** Involved in conceptualization, reviewed and edited the manuscript, supervised data acquisition and provided funding.

## Data Availability Statement

All data supporting the findings of this study have been deposited in Zenodo and are publicly available at: https://doi.org/10.5281/zenodo.19068620

